# Mature adipocytes inhibit differentiation of myogenic cells but stimulate proliferation of fibro-adipogenic precursors derived from trout muscle *in vitro*

**DOI:** 10.1101/2024.05.15.594377

**Authors:** Valentine Goffette, Nathalie Sabin, Jerôme Bugeon, Sabrina Jagot, Isabelle Hue, Jean-Charles Gabillard

**Affiliations:** INRAE, LPGP, 35000, Rennes, France

**Author notes:** Corresponding author: INRAE, Laboratoire de Physiologie et Génomique des Poissons, Campus de Beaulieu, 35042 Rennes cedex, France. This research was funded, in whole or in part, by ANR, Grant #ANR-20-CE20-0013-01. A CC-BY public copyright license has been applied by the authors to the present document and will be applied to all subsequent versions up to the Author Accepted Manuscript arising from this submission, in accordance with the grant’s open access conditions.

**Keywords:** satellite cells, adipose tissue, pax7, pdgfra, myogenin, co-culture

## Abstract

Interactions between tissues and cell types, mediated by cytokines or direct cell-cell exchanges, regulate growth. To determine whether mature adipocytes influence the *in vitro* development of trout mononucleated muscle cells, we developed an indirect coculture system, and showed that adipocytes (5×10^6^ cells/well) derived from perivisceral adipose tissue increased the proliferation (BrdU^+^) of the mononucleated muscle cells (26% *versus* 39%; P<0.001) while inhibiting myogenic differentiation (myosin^+^) (25% *versus* 15%; P<0.001). Similar effects were obtained with subcutaneous adipose tissue-derived adipocytes, although requiring more adipocytes (3×10^7^ cells/well *versus* 5×10^6^ cells/well). Conditioned media recapitulated these effects, stimulating proliferation (31% *versus* 39%; p<0.001) and inhibiting myogenic differentiation (32% *versus* 23%; p<0.001). Adipocytes began to reduce differentiation after 24 hours, whereas proliferation stimulation was observed after 48 hours. While adipocytes did not change *pax7*^+^ and *myoD1/2*^+^ percentages, they reduced *myogenin*^+^ cells showing inhibition from early differentiation stage. Finally, adipocytes increased BrdU^+^ cells in the *Pdgfrα*^+^ population but not in the *myoD*^+^ one. Collectively, our results demonstrate that trout adipocytes promote fibro-adipocyte precursor proliferation while inhibiting myogenic cells differentiation *in vitro*, suggesting the key role of adipose tissue in regulating fish muscle growth.

## Introduction

The interaction between adipose and muscle tissues has been highlighted in numerous studies in mammals and competition between the growth of both tissues has been reported^1–6^. In the post-embryonic stage, an increase in adipose tissue mass has a negative effect on muscle mass^7^. In fish, evidences for these interactions remain limited. In zebrafish, a defect in muscle development has been shown to increase intramuscular adipocyte infiltration^8^. Genetic selection of rainbow trout (*Oncorhynchus mykiss*) for muscle lipid content, also indicates that higher adiposity is associated with a higher proportion of large muscle fibers and a lower proportion of small fibers^9,10^.

Adipose tissue is mainly composed of mature adipose cells called adipocytes, filled with lipids contained in a unique droplet. In rainbow trout, two main adipose tissue deposits have been identified (perivisceral or subcutaneous)^11^, with specific proteomes and gene expression profiles^11,12^. In fish, the extraction and culture of mature adipocytes (MAs) has been reported in sea bream^13^, tilapia^14^ and trout^15^, but remains difficult to achieve due to the buoyancy of MAs and their limited survival in culture^16,17^. In numerous *in vitro* studies, authors prefer to use adipocytes derived from differentiated fibro-adipogenic precursors (FAP), although there are some differences with primary MAs, such as a smaller droplet size with atypical appearance^18–22^.

Large and fast-growing fish such as trout exhibit continuous growth resulting from fiber hypertrophy and the formation of new muscle fibers known as hyperplasia^23^, at least during the exponential growth phase (up to 1 year). Fiber hypertrophy and hyperplasia require the presence of muscle stem cells, which are called satellite cells due to their localization between the basal lamina and the fiber^24^. Satellite cells are quiescent in normal adult muscle and express *pax7* gene, notably involved in their renewal. Satellite cell activation is rapidly followed by the onset of *myoD* expression, while *myogenin* marks the start of differentiation and *myosin* the end^25^. In fish, protocols have been developed to isolate mononucleated muscle cells (MMCs) from muscle tissue and selectively retain those that adhere to the culture dish for further analysis of their proliferation and myogenic differentiation *in vitro*. Characterization of the extracted MMCs in trout indicates that approximately 60% of the cells are MyoD^+^ and thus are myogenic cells, while the identity of the remaining cells is still unknown^26^. The MMCs have been characterized in mammals, and it’s noteworthy that, in addition to myogenic precursors, a significant proportion of FAPs expressing *pdgfr*α are also found as muscle resident cells^27,28^.

Coculturing of cells from muscle and adipose tissue is now common in mammals for investigating cellular communication. A preliminary study conducted in sheep^29^ demonstrated that coculturing satellite cells with preadipocytes inhibits differentiation of myogenic cells, induces myotube atrophy and insulin resistance^30–32^. Furthermore, when immortalized adipogenic cell lines are cocultured in the presence of muscle cells, preadipocyte differentiation decreases, which is associated with a reduction in lipid accumulation^33^. Over the past decade, it has become clear that adipose and muscle cells communicate through multiple secreted factors, known as adipokines and myokines^34–36^, respectively. In mammals, adiponectin, a key adipokine, play a critical role in lipid metabolism^37^ and stimulates glucose uptake and fatty acid oxidation in muscle^38^. Other adipokines, such as leptin, are involved in the development of insulin resistance^39^ in muscle. Among the myokines, myostatin inhibits preadipocyte myogenic differentiation *in vitro*, and limits the formation of lipid deposits^40^. Additionally, muscle-derived interleukin-6 is known to increase uptake and oxidation of fats^41^ as well as adipocyte lipolysis^42,43^.

In fish, our understanding of the mechanisms underlying the interactions between adipose and muscle tissues is very limited. Numerous myokines and adipokines have been identified in fish, including in rainbow trout^45–47^, but there is limited evidence for their role in the cross-talk between these two tissues^44^. For example, in trout, receptors for adiponectin are found in muscle with differential regulation of their expression depending on situations such as fasting, suggesting a possible cross-talk between adipose tissue and muscle^48^.

Although primary cultures of MAs and MMCs from fish have been the subject of monoculture studies regarding their development, co-culture techniques have never been used to study cross-talk between these cell types. The aim of this study was to determine whether mature adipocytes (MAs) influence the *in vitro* development of rainbow trout mononucleated muscle cells (MMCs). Primary mature adipocytes will provide a unique *in vitro* opportunity to better mimic *in vivo* effects on muscle growth. We cocultured these cells in a transwell system to avoid physical cell-cell interactions, but to allow cell-cell communication via soluble molecules, which is particularly relevant given that perivisceral and subcutaneous adipose tissues have no direct contact with skeletal muscle. Comparison of mononucleated muscle cell proliferation and myogenic differentiation in the absence or in presence of adipocytes evidenced a specific cross-talk from adipocytes to fibro-adipogenic progenitors and myogenic cells derived from rainbow trout muscle.

## Materials and Methods

### Animals

Rainbow trout (*Oncorhynchus mykiss*) were reared in a recirculating rearing system at the Fish Physiology and Genomics Laboratory under natural simulated photoperiod and at 12 ± 1 °C. Fish were fed daily *ad libitum* with a commercial diet. Fish were anesthetized with tricaine at 50 mg/L and euthanized with tricaine at 200 mg/l. All the experiments presented in this article were developed in accordance with the current legislation on the ethical treatment and care of laboratory animals (décret no. 2001-464; European directive 2010/63/EU).

### Isolation and culture of mononucleated cells derived from trout muscle

For all studies, mononucleated cells were isolated from the dorsal part of the white muscle of juvenile trout (5 to 30 g body weight) as previously described^26^. Briefly, 20 to 80 g of white muscle were mechanically dissociated and enzymatically digested prior to filtration (100 μm and 40 μm). The cells were seeded onto poly-L-lysine and laminin precoated glass coverslips placed in a 24-well plate at a density of 80,000 cells/cm^2^ and incubated at 18 ^◦^C. Cells were cultured for 2 days in Dulbecco’s modified Eagle’s medium (DMEM) containing 10% fetal bovine serum (FCS) and 1% antibiotic-antimycotic solution. Cells were washed daily with DMEM 1% of antibiotics. The medium was then changed to a 1:1 DMEM and Leibovitz’s L-15 medium containing 10% FCS for last 3 days of monoculture or coculture. Finally, cells were washed twice with PBS and fixed with ethanol/glycine buffer (100% ethanol, 50 mM glycine, pH 2) or fixed in 4% paraformaldehyde for *in situ* hybridization and then preserved in 100% ethanol.

### Isolation and culture of mature adipocytes derived from trout adipose tissue

For all studies, mature adipocytes were isolated from two deposits of adipose tissue: perivisceral (PVA) or dorsal subcutaneous (SA) of trout (150 to 500 g body weight), as previously described^15^. Briefly, 5 to 40 g of adipose tissue were collected, cut into thin pieces, and incubated for 90 min in Krebs – Hepes buffer containing collagenase type II (125 U/ml; Sigma C6885) and 1 % BSA in a shaking platform at 17 °C. The cell suspension was then filtered at 300 µm for perivisceral tissue and 200 µm for subcutaneous tissue. After two washes by flotation in Krebs-Hepes 1 % BSA and two washes by flotation in Krebs-Hepes 2% BSA, cells were counted and cultured in DMEM/L15 (1:1) 10% FCS and 1% antibiotic-antimycotic solution, directly in a transwell (Corning 3413) at a number of 5×10^6^, unless another concentration is specified, until monoculture or coculture.

### Adipocyte size distribution

At the end of cell extraction, a portion of the cells was placed between a microscope slide and a coverslip for microscopic imaging (Nikon digital camera coupled to an Olympus IX70 microscope). Ten images were captured per preparation. A Fiji macro was then used to automatically measure the diameter of each adipocyte. To avoid considering free lipid droplets, data points below 10 µm were excluded from the analyses.

### Coculture of mononucleated muscle cells and mature adipocytes

After extraction, mononucleated muscle cells were cultured, at the density of 80,000 cells/cm2 in DMEM containing 10% FCS until day 2. After extraction, 5×10^6^ adipocytes were cultured directly in a transwell (0.33cm^2^) with a 0.4 µm porous membrane in DMEM/L15 (1:1) with 10% FCS for 1 day. On day 2 of the muscle cell culture, transwells containing the mature adipocytes (24h) were placed on top of wells containing MMC for 72h of coculture (day 2 to day 5) in DMEM/L15 (1:1) with 10% FCS. Finally, the MMCs were washed twice with PBS and fixed with ethanol/glycine buffer (100% ethanol, 50 mM glycine, pH 2) or fixed in 4% paraformaldehyde and then preserved in 100% ethanol.

### Preparation of conditioned medium (CM)

As illustrated in Figure 3a, medium was collected at day 5 (72h of coculture) to obtain coculture conditioned medium (CM CC). Mature adipocytes and MMC were cultured separately in DMEM/L15 (1:1) 10% FCS until day 5 to obtain monoculture conditioned medium (CM MMC, CM MA). All conditioned media were frozen at −80 °C after collection.

### Analysis of proliferation and myogenic differentiation of mononucleated muscle cells

Cells were cultured in presence of 10 μM BrdU during 24H before fixation at day 5. The cells were fixed with ethanol/glycine buffer (100% ethanol, 50 mM glycine, pH 2). After three washes PBS, MMC were saturated for 1 h with 3% BSA, 0.1% Tween-20 in PBS (PBST). Cells were incubated for 30 min at 37 °C with mouse anti-BrdU (11296736, Roche. 1/10) then washed before incubation at room temperature for 3 h with the primary antibody anti-myosin heavy chain (MF20. 1/50; Hybridoma Bank). Finally, cells were incubated with two secondary antibody anti-mouse for 1 hour (anti-mouse IgG1 Alexa 488 A21121, anti-mouse IgG2b Alexa 594 A21145, Fisher 1/1000). Nuclei were stained with a solution of 0.1 µg/mL DAPI (D8417, Sigma) in PBS applied to the cells for 5 min. Cells were then mounted in Mowiol and photographed using a Nikon digital camera coupled to a Nikon Eclipse 90i microscope. Five images were taken per well and the number of BrdU positive nuclei, the number of nuclei in the myosin positive cells and the total number of nuclei was automatically calculated using FIJI software^49^.

### RNAscope *in situ* hybridization and BrdU detection

Detection by *in situ* hybridization of *pax7*, *myoD1/2*, *myogenin* and *pdgfrα* transcripts in fixed MMC was performed as previously described^50^. Briefly, MMCs were fixed with 4 % PFA overnight at 4°C and stored in 100 % (v/v) ethanol at −20 °C until use. Hybridization was performed using the RNAscope Multiplex Fluorescent Assay v2 (Bio-Techne #323100) according to the manufacturer’s protocol. After rehydration, cells were placed in hydrogen peroxide solution (Bio-Techne #322335) for 10 minutes, followed by Protease III solution (1:15) (#322337; Bio-Techne) at 40°C for 10 minutes. Due to the presence of two *myoD* genes in the trout genome, we designed a set of probes targeting MyoD1 and MyoD2 mRNA. This probe set, as other probes, was hybridized at 40°C for 2 hours. The *pax7*, *myoD1/2* (condition 72h), *myogenin* or *pdgfr*α transcripts were detected using the fluorescent dyes Opal 520 (#OP-001001, Akoya Biosciences) and *myoD1/2* (condition 24h) was detected using the fluorescent dyes Opal 620 (#OP-001004, Akoya Biosciences). For the cells under the 72h condition, after two washes with PBS, proliferation staining was followed by in situ labeling. Cells were saturated with 3% BSA in 0.1% Tween-20 in PBS (PBST) for 1 hour. Cells were incubated with rabbit anti-BrdU (#PA5-32256, 1/750) for 4 hours at room temperature, washed, and then incubated with anti-rabbit the secondary antibody (anti-rabbit IgG Alexa 594 A21122, Fisher 1/1000) for 1 hour at room temperature. Cell nuclei of all conditions were stained with a solution of 0.1 µg/mL DAPI (D8417, Sigma) in PBS applied to the cells for 5 minutes. The cells were then mounted in Mowiol and photographed using a Nikon digital camera coupled to a Nikon Eclipse 90i microscope. Five images were taken per well and 4 to 6 wells were used per condition.

### Automated quantification of cells labeled by *in situ* hybridization

To automatically quantify the number of cells expressing these gene, we adapted a macro-command in the Fiji software to quantify puncta corresponding to the RNAscope labeling, per cell^50^. A cell was considered positive if at least 5 puncta were detected in a cell. Our quantification method is available at https://gitlab.univ-nantes.fr/SJagot/fijimacro_rnascopecells.

### Statistical analyses

For analyses comparing proliferation or differentiation across multiple conditions and experimental repetitions (Fig. 1,2,3,4), statistical analyses were conducted using the following approach: when sample size (n) exceeded 10 and the assumptions of parametric tests (such as normal distribution and homogeneity of variances) were met (confirmed by Shapiro-Wilk and Levene tests, respectively), a two-factor ANOVA followed by Tukey post hoc tests was applied. Otherwise, the non-parametric Scheirer-Ray-Hare test followed by Dunn’s post hoc test with Bonferroni correction was utilized. The p-values reported in the text correspond to comparisons between different experimental conditions. The comparison of mean adipocyte diameters (n>10 and meeting the test conditions validated by Shapiro-Wilk and Levene tests) was conducted using a t-test. Statistical analysis was performed using the chi-square test to compare the proportions of adipocytes with a diameter greater than 25µm between the two extraction in figure 2. The comparison of our two groups with sample sizes below 10 was conducted using the non-parametric Wilcoxon test. Specifically, the test was applied to compare the expression of genes between MMC and MMC + MA (Fig. 5,6). All statistical tests were two-tailed, and the significance level (alpha) was set at 0.05. All the statistical analyses were performed with R (version 4.1.3)

## Results

### Mature adipocytes influence the growth of mononucleated muscle cells *in vitro* in a dose-dependent manner

To determine whether mature adipocytes (MAs) could influence the *in vitro* growth of mononucleated muscle cells (MMCs), we measured the proliferation (BrdU^+^) and the myogenic differentiation (myosin^+^) of MMCs by immunofluorescence in the presence or absence of MA extracted from perivisceral adipose tissue (Fig. 1a). After 72 hours of coculture or MMC monoculture, results showed the presence of numerous BrdU^+^ nuclei as well as mononucleated and multinucleated (myotubes) cells expressing myosin (Fig. 1b, c). Measurement of the percentage of BrdU^+^ nuclei showed a significant (P<0.001) increase in the proliferation of MMCs in the presence of MAs (5×10^6^), with approximately 39% of cells having proliferated in the last 24 hours, compared to approximately 26% in the absence of MAs (Fig. 1d). In contrast, the percentage of nuclei expressing a late myogenic differentiation marker, myosin, decreased in the presence of MAs (5×10^6^) compared to MMC monoculture (25% *versus* 15%; P<0.001)(Fig. 1e).

**Figure 1.**
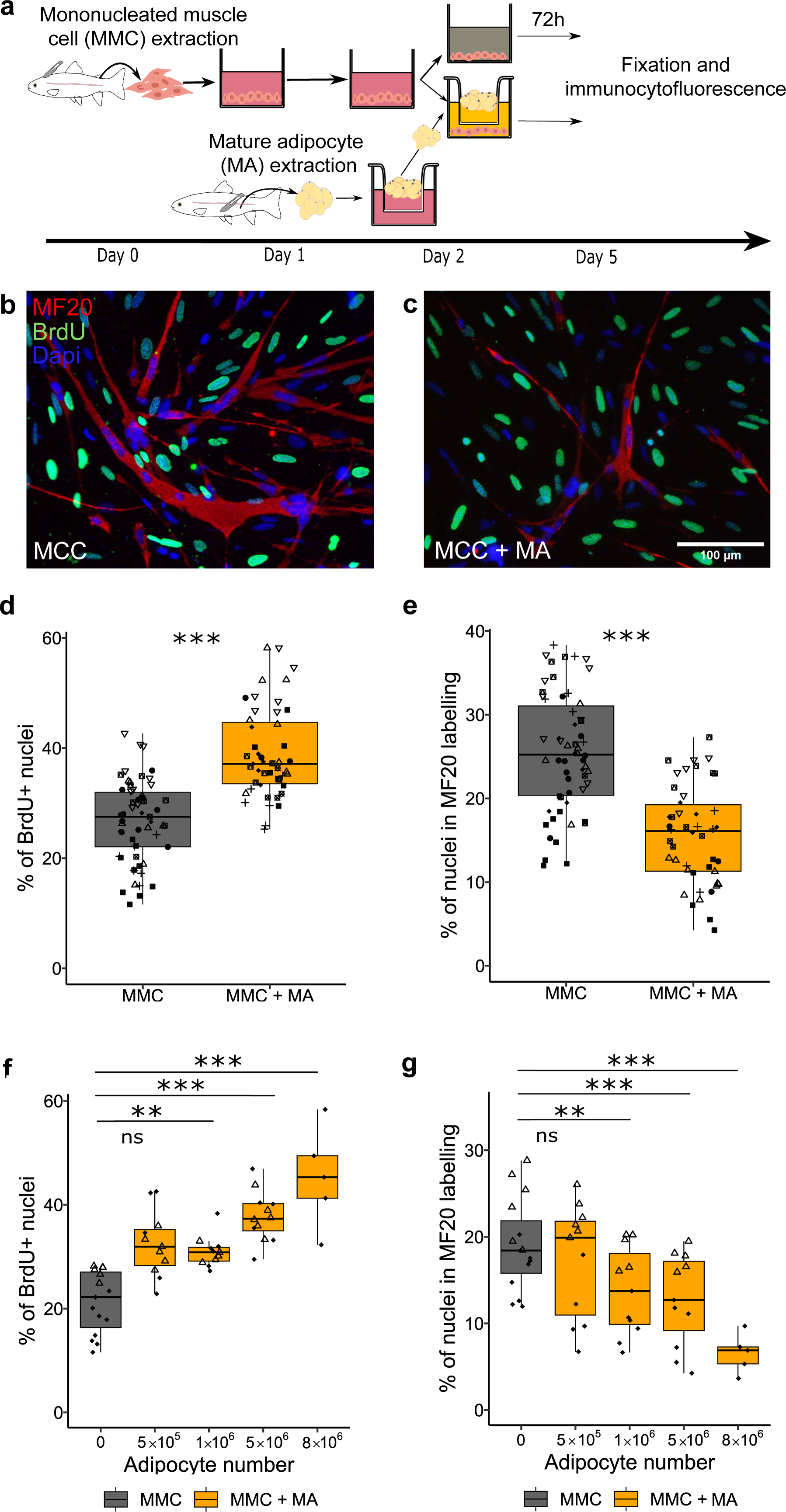
Mature adipocytes stimulate proliferation but inhibit differentiation of mononucleated muscle cells. (a) Diagram of the isolation, extraction, culture and co-culture of mononucleated muscle cells (MMC) and mature adipocytes (MA) from perivisceral adipose tissue of rainbow trout. (b, c) Images of immunocytofluorescence analysis for myosin (MF20, red), BrdU (green) in MMC alone (MMC) or cocultured with MA (MMC+MA) after fixation on day 5. Cell nuclei are stained with Dapi (blue). (d, f) Quantification of cell proliferation, measured by BrdU incorporation, and myogenic differentiation (e, g), measured by immunocytofluorescence labeling of myosin (MF20). Compared to monocultured MMCs, MMCs cocultured with MA for 72 hours display higher proliferation (d) and lower differentiation (e). (f, g) Dose-dependent effect of adipocyte number (0, 5×10^5^, 1×10^6^, 5×10^6^, 8×10^6^) on proliferation (BrdU) and differentiation (MF20) of MMCs. The different symbols correspond to different experiments. Statistical significance was determined by two-factor ANOVA followed by Tukey post hoc tests, or Scheirer-Ray-Hare tests with Dunn post hoc and Bonferroni correction. Significance levels: ns (not significant), ** (p < 0.01), *** (p < 0.001).

After observing the effect at a given number of MAs, we investigated the possibility of a dose-dependent effect. Using different amounts of MAs (5×10^5^ up to 8×10^6^), we observed a clear positive dose-dependent effect on the proliferation of MMCs (Fig. 1f), together with a negative dose-dependent effect on the myogenic differentiation (Fig. 1g). These results indicated that even as few as 1×10^6^ MAs were sufficient to affect both the proliferation and the myogenic differentiation of MMCs.

### Adipose tissue origin influences the size of mature adipocytes and their effect on mononucleated muscle cells

To characterize the MAs that have been extracted from perivisceral and subcutaneous adipose tissue of rainbow trout, analysis of bright field images (Fig. 2a) showed a lower mean diameter in the subcutaneous (SA) compared to the perivisceral (PVA) extraction (21.5 µm *versus* 24.3 µm, P<0.001) (Fig. 2b). Overall, we observed a lower proportion of MAs >25 µm in SA compared to PVA (25% *versus* 35%, P<0.001) (Fig. 2c).

**Figure 2.**
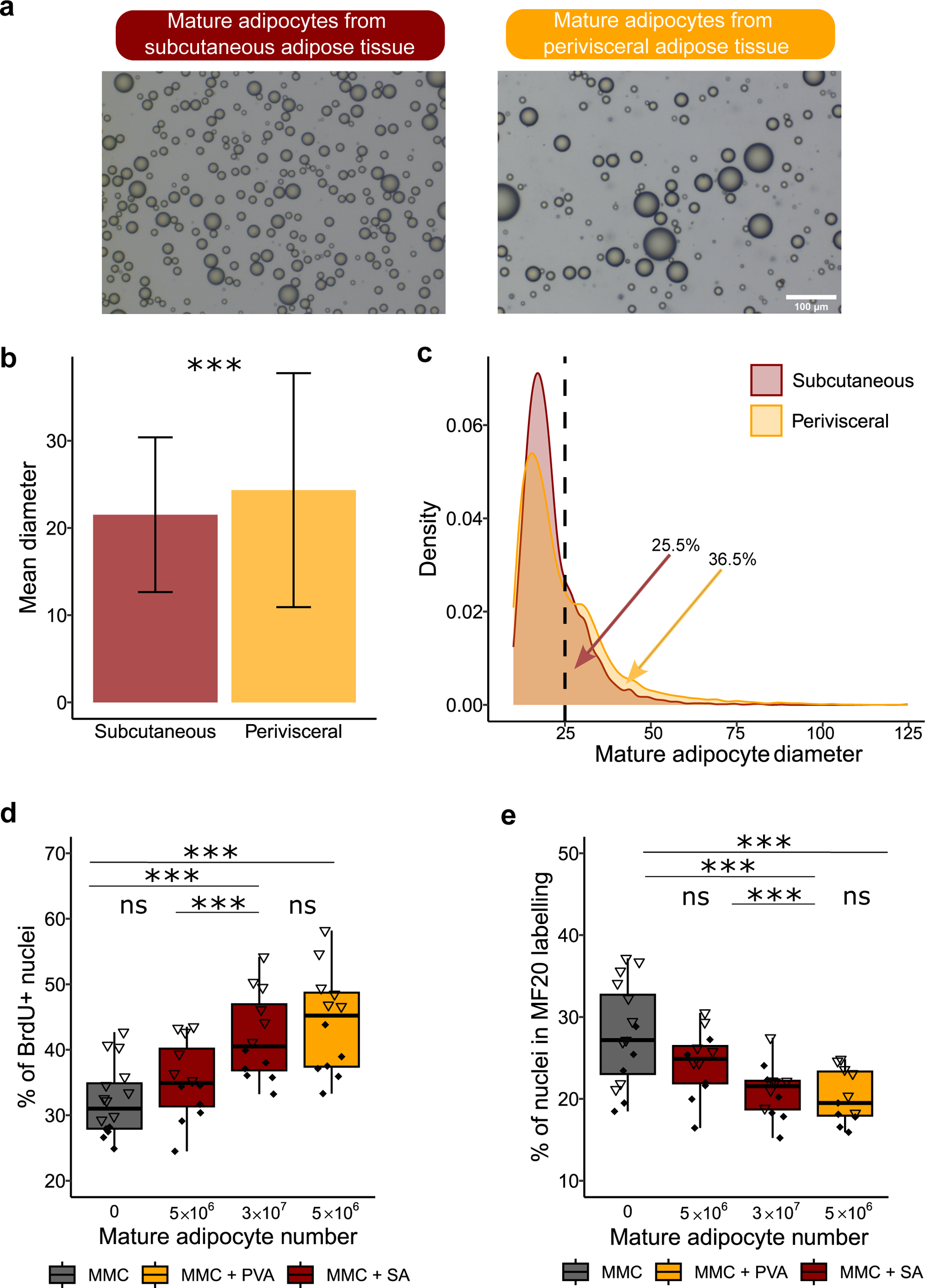
Comparison of the effect of mature subcutaneous and perivisceral adipocytes on MMC. (a) Bright field images of mature adipocytes (MA) extracted from subcutaneous (SA) (left, red) and perivisceral (PVA) (right, orange) adipose tissue. (b) Comparison of the mean adipocyte diameter (>10µm) between the two extractions, showed a significantly higher adipocyte diameter in PVA. (c) Distribution of adipocyte size (>10µm) from SA (red) and PVA (orange) showed a higher proportion of adipocytes with a diameter >25 µm in PVA compared to those from SA. (d) MMC proliferation, measured by BrdU quantification and (e) myogenic cell differentiation, measured by immunocytofluorescence labeling of myosin (MF20). Compared to MCC monoculture, MMC cocultured for 72 hours with MA extracted from perivisceral adipose tissue (MMC + PVA), displayed higher proliferation and lower differentiation. When MMCs were cocultured with MA extracted from subcutaneous adipose tissue (MMC + SA) at an equivalent cell number (5×10^6^) and duration (72 h), no significant differences were observed compared to MMCs cultured alone. However, an increase to 3×10^7^ cells showed a similar effect on both proliferation and differentiation parameters, suggesting a response dependent on cell quantity. The different symbols correspond to different experiments. Statistical significance was determined by two-factor ANOVA followed by Tukey post hoc tests, or Scheirer-Ray-Hare tests with Dunn post hoc and Bonferroni correction. Significance levels: ns (not significant), * (p < 0.05), ** (p < 0.01), *** (p< 0.001).

We wondered if MAs extracted from subcutaneous adipose tissue would have the same effects on MMCs as previously observed with perivisceral MAs. To address this question, we established cocultures with MMCs and different amounts of subcutaneous MAs. For a same amount of MAs added (5×10^6^), the percentage of MMC proliferation did not show a significant difference in the presence or absence of subcutaneous MAs (32% *versus* 35%; p = 0.13), whereas a clear increase is observed with perivisceral MAs (32 *versus* 44%; P<0.001) (Fig. 2d). When examining the percentage of nuclei in myosin^+^ cells, no significant reduction in myogenic differentiation was observed in MMCs cocultured with 5×10^6^ subcutaneous MAs compared to monoculture of MMCs (25% *versus* 28%; p = 0.8). In contrast, with the same amount (5×10^6^) of perivisceral MAs, a significant decrease in the percentage of nuclei in myosin^+^ cells was observed (28% *versus* 20%; p =0.002) (Fig. 2e). However, a 6-fold increase (3×10^7^) in the number of subcutaneous MAs gave results comparable to 5×10^6^ perivisceral MAs, i.e. an increase in proliferation (32% *versus* 42%; P<0.001) (Fig. 2b) and a decrease in myogenic differentiation (28% *versus* 21%; p = 0.008) (Fig. 2e) compared to MMC monoculture.

### Mature adipocyte-derived soluble factor(s) influence the *in vitro* development of mononucleated muscle cells

In our previous experiments, we used indirect cocultures, in which both cell types share a common culture medium but are physically separated by a porous membrane (0.4 µm). The factor(s) contributing to the observed effects on MMCs should be soluble, smaller than the transwell’s pores, and able to diffuse through the culture medium. In order to confirm this hypothesis, we cultivated MMCs with different conditioned media as shown in Figure 3a. Our analyses showed no significant effect on both proliferation (31% *versus* 32%; p = 0.99) (Fig. 3b) and myogenic differentiation (32% *versus* 31%; p = 0.88) (Fig. 3c) of MMCs cultured with medium conditioned by a previous MMC culture (CM MMC) alone compared to fresh monoculture of MMCs (MCC). In contrast, medium conditioned by either coculture (CM CC) or by MAs alone (CM MA) increased proliferation (31% *versus* 38%; P<0.001, 31% *versus* 39%; P<0.001) (Fig. 3b) and decreased myogenic differentiation (32% *versus* 25%; P<0.001, 32% *versus* 23%; P<0.001) (Fig. 3c) of MMCs. This effect of conditioned medium was comparable to that obtained with freshly extracted adipocytes cells (proliferation: 31% *versus* 40% P<0.001, myogenic differentiation: 32% *versus* 19%; P<0.001) (Fig. 3b,c). We observed the same effect with a medium conditioned by coculture of adipocytes and MMCs (CM CC) as with mature adipocytes alone (CM MA).

**Figure 3.**
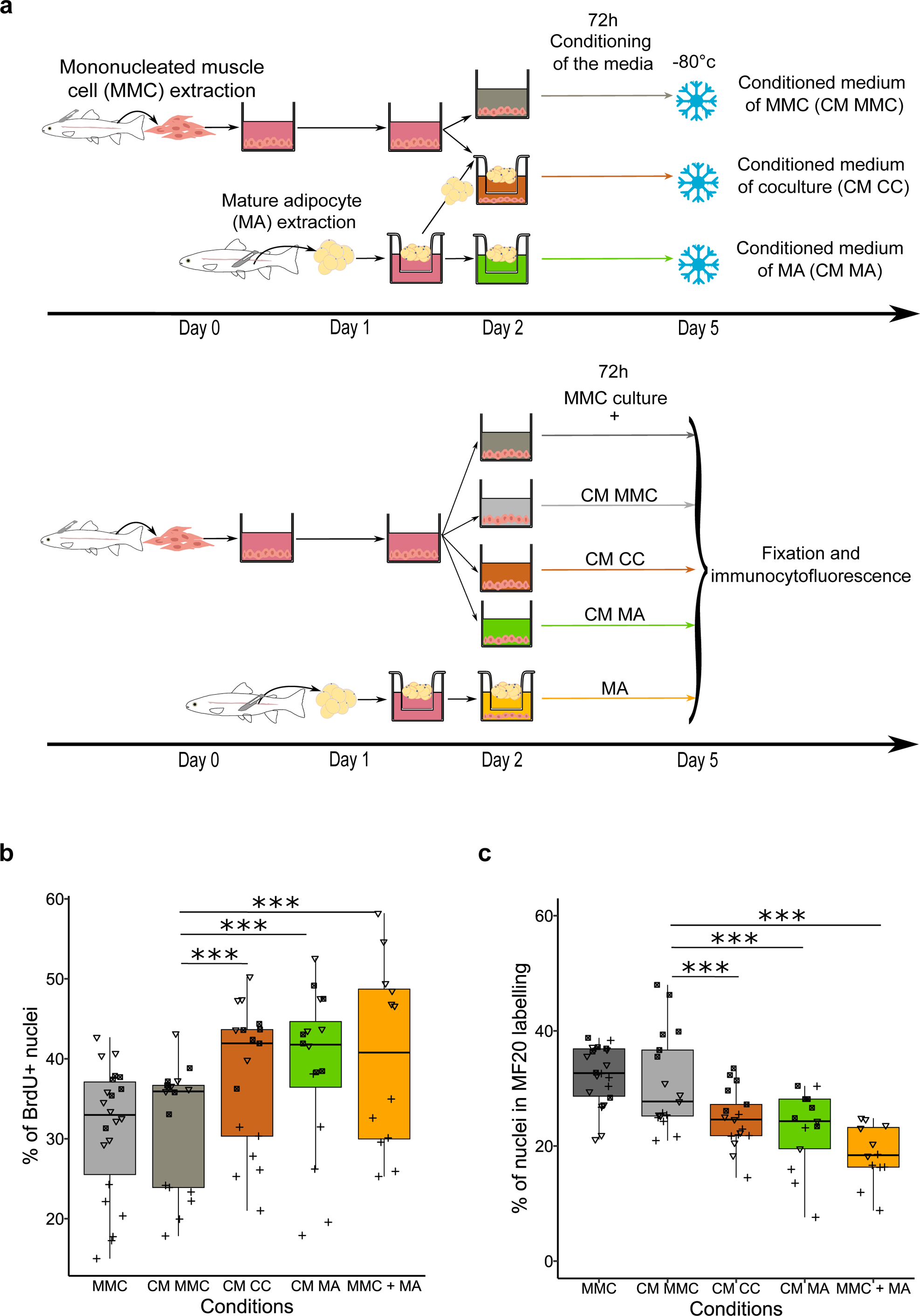
Conditioned medium with mature adipocytes alone is sufficient to influence the development of mononucleated muscle cells *in vitro*. (a) Diagram of the isolation, extraction, culture for media conditioning, and diagram of isolation, extraction, and culture of mononucleated muscle cells (MMC) with either mature adipocytes (MMC + MA) or conditioned media: conditioned medium with MMC alone (CM MMC), conditioned medium with coculture (CM CC), conditioned medium with mature adipocytes (CM MA). (b) MMC proliferation, measured by BrdU quantification. Compared to MMC monoculture (MMC) or MMC culture with conditioned medium with MMC (CM MMC), exposure to conditioned medium with mature adipocytes (CM MA) or with coculture (CM CC) led to an increase in MMC proliferation (p<0.001). This effect mirrors that observed in the presence of freshly extracted mature adipocytes from perivisceral tissue (MMC + MA). (c) Opposite effects were observed on myogenic cell differentiation, as revealed by immunocytofluorescence labeling of myosin (MF20). No significant change was observed with CM MMC, whereas CM MA and CM CC showed a marked reduction, similar to the effect of MA. The different symbols correspond to different experiments. Statistical significance was determined by two-factor ANOVA followed by Tukey post hoc tests, or Scheirer-Ray-Hare tests with Dunn post hoc and Bonferroni correction. Significance levels: ns (not significant), * (p < 0.05), ** (p < 0.01), *** (p < 0.001).

### Mature adipocytes inhibit early differentiation of myogenic cells *in vitro*

To investigate the dynamics of the interaction between MAs and MMC development *in vitro*, we established coculture kinetics to determine the time required to observe a significant effect on proliferation and on myogenic differentiation (Fig. 4a). While a clear increase in proliferation of MMCs was observed after 48 h (29% *versus* 37%; P<0.001) and 72 h (30% *versus* 39%; P<0.001) of coculture, 24 h of exposure was not sufficient to observe a significant effect on MAs (19% *versus* 23%; p = 0.051) (Fig. 4b). In contrast, a small but significant decrease in myogenic differentiation was observed already after 24 h of adipocyte exposure (9% *versus* 5%; p =0.0157), and was further enhanced at 48 h and 72 h (Fig. 4c).

**Figure 4.**
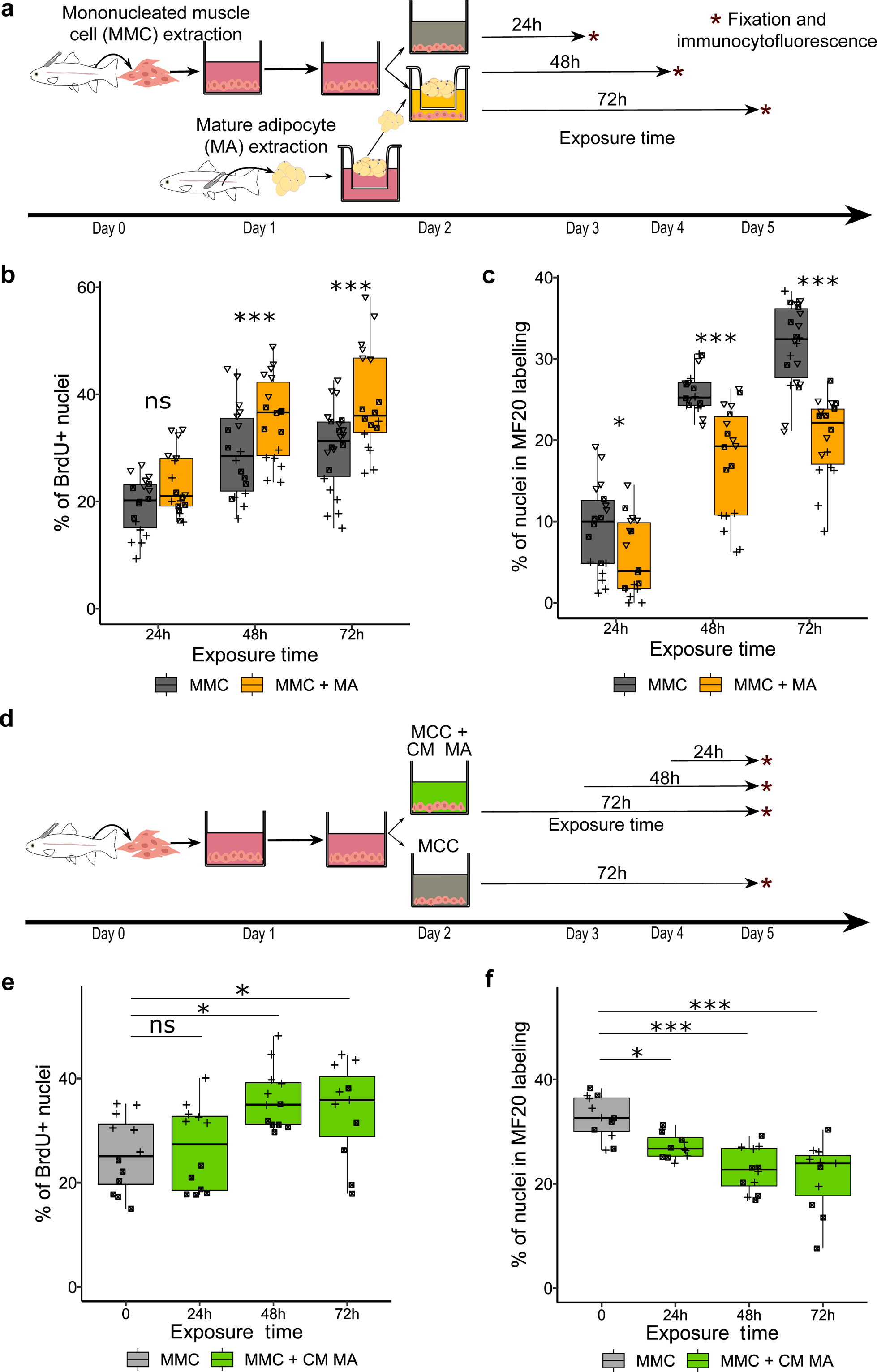
Time-dependent effects of mature adipocytes on the development of mononucleated muscle cells *in vitro*. (a) Diagram of the isolation, extraction and coculture of mononucleated muscle cells (MMC) and mature adipocytes (MA) from perivisceral adipose tissue, with fixation of MMC at different time points after the onset of coculture (24 h, 48 h, 72 h). (b) MMC proliferation, measured by BrdU quantification, showed no significant difference at 24 h in the presence of MA, whereas a marked increase in proliferation was observed at 48 h and 72 h of coculture. (c) Myogenic cell differentiation, measured by immunocytofluorescence labeling of myosin (MF20), was slightly decreased at 24h in the presence of MA, followed by a pronounced decrease at 48h and 72h of coculture. (d) Diagram of the isolation, extraction and culture of MMCs with adipocyte-conditioned medium (CM MA) using different exposure times (24 h, 48 h, 72 h) but fixation on day 5 for each condition. (e) MMC proliferation, measured by BrdU quantification, showed that a 24 h exposure to conditioned medium between day 4 and day 5 is not sufficient to increase proliferation (p=0.3), in contrast to 48 h or 72 h exposure to conditioned medium. (f) Myogenic cell differentiation, measured by immunocytofluorescence labeling of myosin (MF20), showed a significant reduction in differentiation already after 24h of exposure to conditioned medium. The different symbols correspond to different experiments. Statistical significance was determined by two-factor ANOVA followed by Tukey post hoc tests, or Scheirer-Ray-Hare tests with Dunn post hoc and Bonferroni correction. Significance levels: ns (not significant), * (p < 0.05), ** (p < 0.01), *** (p < 0.001).

To determine whether the responsiveness of MMCs to adipocyte-derived factor(s) changes during the culture, we exposed the MMCs to conditioned medium for different time periods and fixed the cells at the same developmental stage (day 5) (Fig4. d). Exposure to conditioned medium for the last 48 hours and the last 72 hours, increased proliferation (26% *versus* 36%; p=0.014, 26% *versus* 34% p=0.033)(Fig. 4e) as well as decreased myogenic differentiation (33% *versus* 23%; P<0.001, 33% *versus* 21%; P<0.001) (Fig. 4f). However, while we did not observe a significant difference in proliferation of MMCs cultured in conditioned medium during the last 24 hours (26% *versus* 27%; p=0.3) (Fig. 4e), we still found a reduction in myogenic differentiation (33% *versus* 27%; p=0.022) (Fig. 4f).

To determine at which stages adipocyte-derived factor(s) inhibits the myogenic program, we performed *in situ* hybridization with markers of satellite cells (*pax7*), myoblasts (*myoD1/2*) and myocytes (*myogenin*), after 24h of coculture (Fig. 5a). As shown in Figure 5b, 24h of coculture did not change the percentage of *pax7*^+^ (73% versus 70%; p=0.55) or the percentage of *myoD1/2*^+^ cells (52% versus 48%; p=0.41). In contrast, we observed a significant decrease in the percentage of *myogenin*^+^ cells after 24 h of coculture compared to monoculture of MMCs (44% versus 38%; P<0.001).

**Figure 5.**
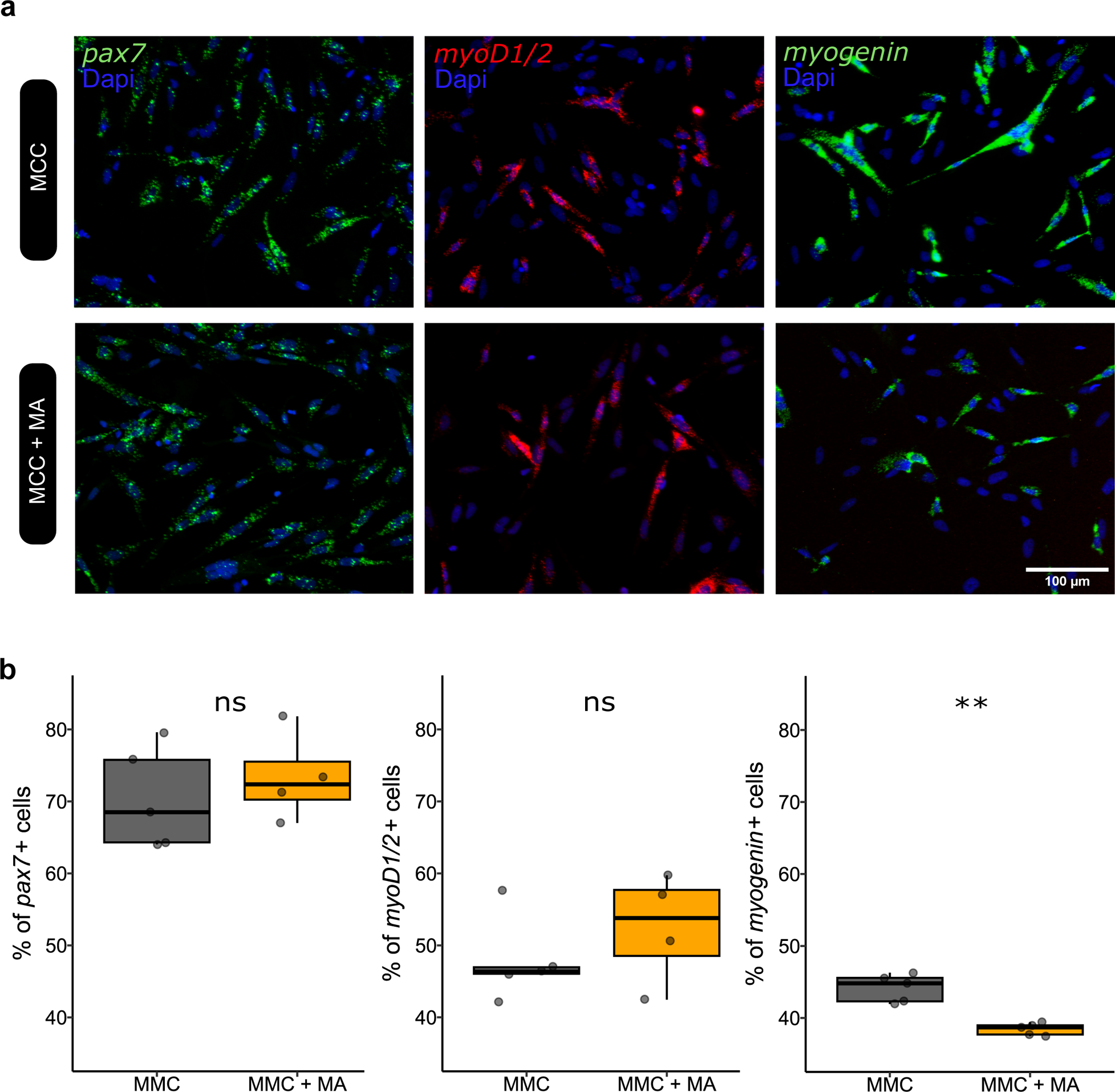
Early inhibition of myogenic differentiation of mononucleated muscle cells by mature adipocytes *in vitro*. (a) Images showing *in situ* hybridization analyses of *pax7* (green) or *myod1/2* (red) or *myogenin* (green) expression on monocultured mononucleated muscle cells (MMC) or MMC cocultured for 24h with mature adipocytes (MA) from perivisceral adipose tissue. Cell nuclei are stained with Dapi (blue). (b) Quantification of MMC percentage expressing *pax7, myod1/2* and *myogenin* revealed that matured adipocytes significantly reduced only the percentage of *myogenin*^+^ cells. Statistical significance was determined by the Wilcoxon test. Significance levels: ns (not significant), * (p < 0.05), ** (p < 0.01).

### Mature adipocytes stimulate proliferation of fibro-adipogenic progenitors but not of myogenic cells *in vitro*

We aimed to further characterize which population of MMCs proliferates in response to adipocyte-derived soluble factor(s). First, we performed *in situ* hybridization on the monoculture of MMCs with markers of fibro-adipogenic progenitors (FAPs) and myogenic cells, i.e. *pdgfrα* and *myoD1/2,* respectively (Fig. 6a). Our results showed that MMC monoculture contained 53% of *myoD1/2*^+^ cells and 55% of *pdgfrα* ^+^ cells, indicating that they represent the two major populations of mononucleated cells derived from white muscle (Fig. 6b). After 72 h of coculture, we observed that the percentage of *pdgfrα* ^+^ cells was increased compared to monoculture (55% *versus* 66%, p = 0.0043), whereas no significant difference was observed for the percentage of *myoD1/2*^+^ cells (53% versus 52%, p=1) (Fig. 6b). To better determine which cell population was stimulated by MAs, we also performed double labeling with BrdU and *in situ* hybridization for *myoD1/2* or *pdgfrα*. The results indicated that the percentage of BrdU^+^ cells within the *myoD1/2*^+^ population was similar between coculture and monoculture conditions (19% versus 19%, p=0.84), whereas the percentage of proliferative cells (BrdU^+^) in the *pdgfrα* ^+^ population increased (24% versus 32%; p = 0.041) (Fig. 6c) when MMCs were cultured in the presence of MAs.

**Figure 6.**
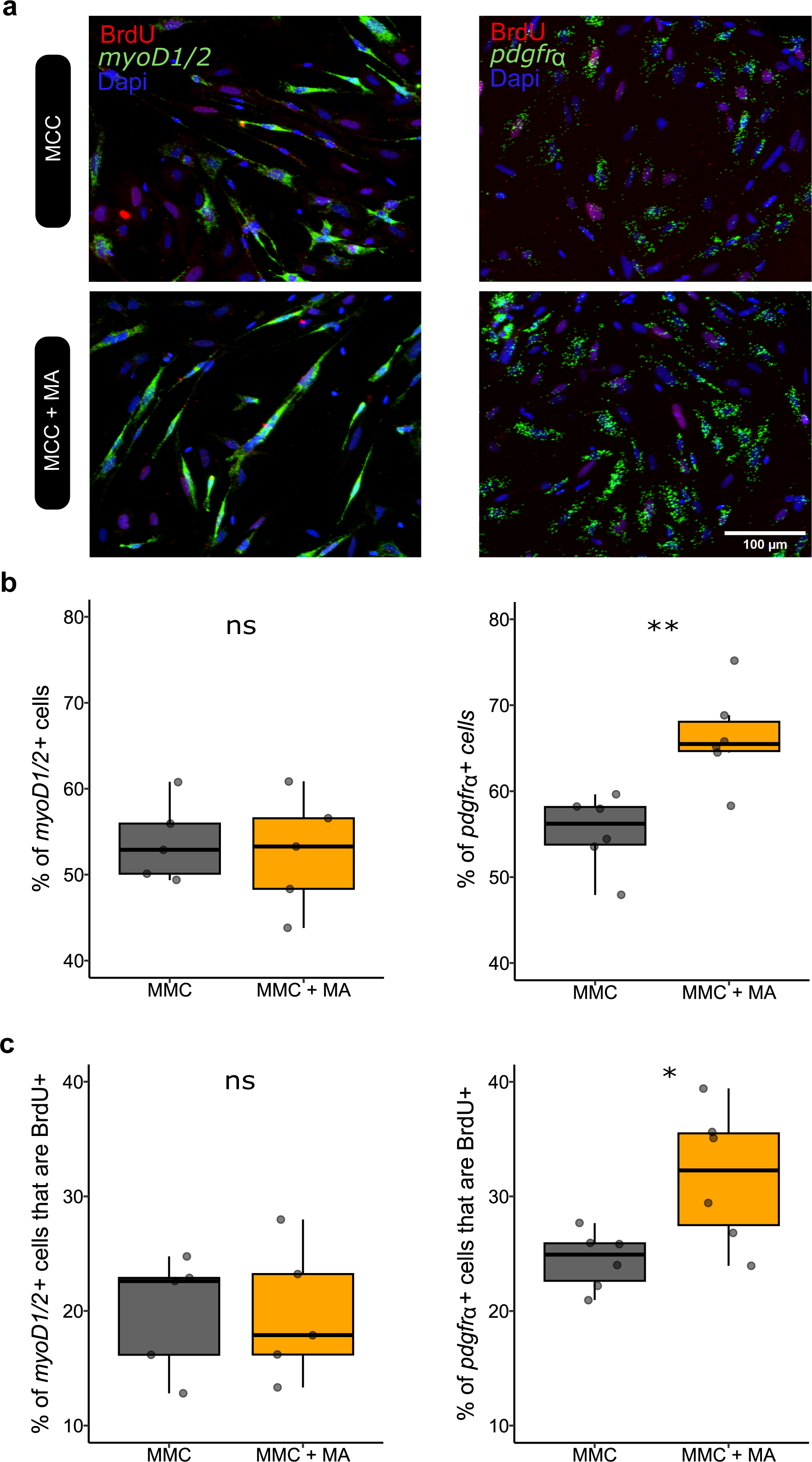
Mature adipocytes stimulate *in vitro* proliferation of fibro-adipogenic progenitors but not of myogenic cells derived from mononucleated muscle cells *in vitro*. (a) Images showing immunocytofluorescence detection of BrdU (red) associated with *in situ* hybridization analysis of *myod1/2* (green) or *pdgfrα* (green) expression on monocultured mononucleated muscle cells (MMC) or MMC cocultured for 72 h with mature adipocytes (MA) from perivisceral adipose tissue. Cell nuclei are stained with Dapi (blue). (b) Quantification of MMC percentage expressing *myod1/2* and *pdgfrα* indicated that mature adipocytes significantly increased *pdgfrα*^+^ cell proportion. (c) Proliferation of each cell type (*myod1/2*^+^ or *pdgfrα*^+^), was analyzed by detection of BrdU incorporation and showed no increase of BrdU^+^ percentage in *myod1/2*^+^ cells in coculture, but a significant increase in proliferation of *pdgfrα*^+^. Statistical significance was determined by the Wilcoxon test. Significance levels: ns (no significance), * (p < 0.05), ** (p < 0.01).

## Discussion

The rainbow trout (*Oncorhynchus mykiss*) is an interesting model to study the communication between adipose and muscle tissues due to its different growth patterns compared to mammals. Indeed, trout exhibit an exponential muscle growth during the post-larval phase, associated with strong hyperplasic and hypertrophic muscle activity. In the specific context of salmonid models, the influence of adipose tissue on muscle growth remains poorly characterized. The aim of this study was to determine whether mature adipocytes from different adipose tissues can influence the proliferation and the myogenic differentiation of mononucleated muscle cells (MMCs) *in vitro*. Our main results provide direct evidence for the existence of cellular communication between mature adipocytes (MAs), fibro-adipogenic progenitors (FAPs) and myogenic cells in trout.

In vertebrates, two preferred storage sites for adipose tissue have been identified, perivisceral and subcutaneous, which are known to have different mobilization and metabolism^51–53^. In trout, differences in the size distribution of MAs and the abundance of certain proteins have been observed between the two tissue types, indicating different metabolic activities ^11,12^. Our results confirm the difference in MA size between both adipose tissues, with a higher proportion of larger MAs in visceral adipose tissue compared to subcutaneous tissue. Because of these differences, we also compared the effect of MAs from both adipose tissues on MMC growth.

To assess the influence of trout MAs on the *in vitro* growth of MMCs, we measured the proliferation and myogenic differentiation in the presence or absence of adipocytes in an indirect coculture system. We used mature primary adipocytes, which brings us closer to *in vivo* conditions, whereas studies in mammals typically use *in vitro* differentiated preadipocytes, which may exhibit different properties. Under our experimental conditions, we observed a strong stimulation of MMCs proliferation by perivisceral mature adipocytes, in a dose-dependent manner (from 1×10^6^ to 8×10^6^ adipocytes). Subcutaneous adipocytes induced the same effect, but at a much higher number (3×10^7^), suggesting a difference in secretome between perivisceral and subcutaneous MAs. Such differences are far from being studied in fish species and only partially in humans ^32,54^. Thus far however, our approach, by using BrdU incorporation, provides a specific and accurate measurement of proliferation in indirect coculture systems. These results are consistent with other studies in mammals showing an increase in proliferation of MMCs in response to preadipocytes or adipocytes, using less specific measures such as MTT assay reflecting viability^55^ or by assessing the increase in total cell number^30^. Our results suggest that mature adipocytes from perivisceral tissue may enhance the proliferation of MMCs in trout *in vivo*.

In addition, our *in vitro* results clearly showed that perivisceral MAs inhibited differentiation of myogenic cells in a dose-dependent manner. Again, subcutaneous MAs induced the same effect but at a much higher number (3×10^7^), confirming the difference in secretome between perivisceral and subcutaneous adipocytes. These results are consistent with previous observations in mammalian models showing a decrease in myotube formation during indirect coculture in different models, such as in immortalized cell lines indirect coculture^56^, in rat muscle progenitors with adipogenic cells derived from rat muscle^30^ and in indirect cocultures of dedifferentiated chicken intramuscular adipocytes^57^. Interestingly, the inhibition of myogenic differentiation by MAs was characterized by a decrease in the percentage of *myogenin*^+^ and *myosin*^+^ cells but not of *pax7*^+^ and *myoD1/2*^+^ cells showing that the inhibition occurs from the early stage of differentiation, preventing the formation of myotubes. Accordingly, Takegahara et al. (2014) show that rat MAs decrease the percentage of myosin^+^ cells but not that of MyoD^+^ cells^30^. Inhibition of MMC myogenic differentiation, observed as early as 24 hours, is earlier than previously reported in the literature for an indirect coculture with MAs whereas coculture durations of 2 to 5 days are generally required to observe such an effect ^30,56,57^. Nevertheless, quantitative RT-PCR analyses show that expression of *pax7*, *myoD*, *myogenin* and *myosin*^57^ is reduced as early as 24 hours in presence of chicken intramuscular preadipoctyes. This apparent discrepancy, can arise from existing differences between preadipocytes and mature adipocytes, but also to the technique used. Together, the marked and rapid reduction in myotube formation by MAs is probably due to early inhibition of myogenic differentiation.

Considering the absence of cell-to-cell contact in our experiments, the observed effects on muscle cells should be due to soluble factors. However, we cannot exclude the possibility that MMCs induce the production of factors by adipocytes, which in turn may affect their proliferation and myogenic differentiation. Our results showed that cultured MMCs with medium conditioned with both MMCs and MAs or with MAs alone, stimulated proliferation and inhibited myogenic differentiation to the same extend as freshly isolated adipocytes. These results confirm that MAs secrete one or more soluble factors that directly influenced MMCs growth *in vitro,* and that the production of these factors by adipocytes is independent of the MMCs. The nature of this factor is unknown, but it is known that adipocytes, as many other cells, secrete various molecules such as proteins, lipids, extracellular vesicles, etc. that could stimulate the proliferation of MMCs and inhibit the differentiation of myogenic cells ^30,56,58^.

Since proliferation and myogenic differentiation are mutually exclusive cellular processes, we wondered whether increased proliferation would cause decreased myogenic differentiation, or vice versa, inducing a time lag in the onset of both effects. The kinetic of MMCs proliferation and myogenic differentiation, indicate the effect of MAs on myogenic differentiation was observed as early as 24 h, while the effect on proliferation was not observed until 48 h. Furthermore, incubation of MMCs with adipocyte-conditioned medium during the last 24 h of the culture (from day 4 to day 5) was sufficient to reduce myogenic differentiation but not proliferation. Taken together, these results indicate that the effects of MAs on myogenic proliferation and myogenic differentiation are only slightly time delayed, which cannot directly explain the increased proliferation of MMCs by MAs. We have previously shown that 2 days after MMCs extraction, some cells proliferate while others start to differentiate^26^, demonstrating the presence of cell subtypes at different stages of the myogenic program in the MMCs extracted from trout muscle. Therefore, we wondered whether adipocyte-secreted factors, in addition to inhibiting myogenic differentiation, would also stimulate the proliferation of the cells that are not yet engaged in the myogenic differentiation. Surprisingly, our results show that the proliferation of myogenic cells (*myoD*^+^) does not account for the observed increase in MMCs proliferation induced by MAs in contrast to the proliferation of fibro-adipogenic progenitors (*pdgfrα*^+^) that is stimulated. Several works report that preadipocytes or adipocytes enhance the proliferation of primary culture of MMCs ^55,59^, but the identity of the proliferative cells has never been investigated. In contrast, MAs-induced stimulation of FAP proliferation has previously been observed in FAPs derived from adipose tissue in human, but never in muscle derived FAPs ^60,61^. Thus, FAP proliferation in response to MAs-derived factor appears to be a conserved mechanism regardless of FAP origin.

In conclusion, we have demonstrated a cross-talk between mature adipocytes and mononucleated muscle cells in trout based on adipocytes-derived secreted factor(s) that stimulates proliferation of FAPs but inhibits differentiation of myogenic cells *in vitro*. Despite these findings, much remains to be explored regarding the diverse secretions of adipose tissue in fish, and further studies are needed to determine which specific adipocyte-derived factors may be responsible for the observed effects on mononucleated muscle cells in our experimental context.

## Acknowledgments

We particularly thank C. Duret for trout rearing and C Rallière for technical assistance in RNAscope analyses. This work was supported the ANR FishMuSC (ANR-20-CE20-0013-01). The fellowship of Valentine Goffette was supported by INRA PHASE and the Région Bretagne.

## Author Contributions

IH and J-CG conceptualized the study; VG performed all the laboratory analyses; JB and SJ developed a macro-command on Fiji software to automated quantification (RNAscope, adipocyte size); VG, IH and JCG analyzed and interpreted the data; JCG and IH. acquired funding; VG, IH and JCG drafted and critically reviewed the manuscript. All authors have read and approved the final manuscript.

## Data availability

The datasets used and analysed during the current study are available from the corresponding author on reasonable request.

